# Changes in cell wall biochemistry affect mechanical properties and growth rate of *Arabidopsis* pollen tubes

**DOI:** 10.1101/2021.11.09.467870

**Authors:** Hannes Vogler, Gautam Munglani, Tohnyui Ndinyanka Fabrice, Christian Draeger, Jan T. Burri, Christof Eichenberger, J. Paul Knox, Jean Claude Mollet, Bradley J. Nelson, Hans J. Herrmann, Christoph Ringli, Ueli Grossniklaus

## Abstract

Pollen tubes maintain cell wall integrity as they rapidly grow towards the ovule, yet need to rupture at a precise moment to release the sperm cells. This biomechanical balance is critical for fertilization and relies on the interplay between turgor pressure and cell wall rigidity. How cell wall composition affects its mechanical properties is, however, not well understood. In this study, we combine experimental and simulation techniques to determine key mechanical parameters using *Arabidopsis* cell wall mutants. We integrated cellular force microscopy with a Finite Element Method-based model to predict growth rates of different mutant pollen tubes. The Finite Element Method-based model allowed us to quantify the effects of cell wall mutations on the time-independent turgor pressure and cell wall elasticity, while cellular force microscopy enabled determination of time-dependent viscoelastic properties of the cell wall. This novel approach can be applied across biological systems and advances mechanical studies of cell and tissue morphogenesis.

## Introduction

In all organisms, mechanical forces and physical constraints influence cellular growth and the development of tissues and, hence, present an essential parameter influencing tissue and organ formation (1). Plant morphogenesis is largely regulated by mechanical equilibrium. Controlled cellular growth involves a finely tuned interplay between turgor pressure as the driving force and the constraining action of the cell wall (2, 3), with the rate of expansion being controlled by local changes in cell wall viscosity (4, 5). The importance of a tight control of this interplay is particularly evident in tip-growing cells such as pollen tubes (PTs), which are arguably the fastest-expanding cells on the planet. While a tight control of cell wall integrity is essential during the growth phase, the final step—the release of the sperm cells for successful double fertilization—requires the PT to rupture at the right moment and place. The ability of the cell wall to respond to such different requirements is directly related to the dynamic control of its mechanical properties.

How the mechanical properties of the cell wall depend on its biochemical composition, which dynamically changes in response to internal and external factors, is largely unknown. Thus, to gain deeper insights into the delicate relationship between turgor pressure and the mechanical properties of the cell wall, we need to learn more about the effect of biochemical changes in the cell wall on these parameters. While turgor pressure is a well-studied quantity, the correlation between the cell wall’s composition and its mechanical properties remains poorly understood. Cell walls are complex supramolecular structures of intertwined polysaccharides and structural proteins. The classical model of the cell wall assumes a network of cellulose microfibrils connected by hemicelluloses that is embedded in a gel-like pectin matrix (6) and reinforced by structural proteins, such as the highly glycosylated extensins (7). Together with callose, all these components are also present in the cell wall of PTs (8). The spatially and temporally controlled expansion of the cell wall is facilitated by local modifications of its composition, leading to rheological adjustments that allow the cell to adapt its size and shape to specific requirements (9–11). Thus, information on bulk material behavior under varied conditions is required to understand the cell wall’s role in regulating cell expansion. In particular, capturing the response of the cell wall to forces exerted at both short and long timescales. These distinct loading modalities yield time-independent (instantaneous) and time-dependent (relaxation) properties, respectively, which are essential to quantify the competing forces that maintain mechanical equilibrium and define cell shape. The instantaneous pliability of the cell is best quantified by its apparent stiffness (a measure of perceived stiffness): a combination of turgor pressure, geometric properties (e.g., cell wall thickness), and intrinsic material properties, such as the elastic modulus (aka Young’s modulus, a measure of elasticity). In contrast, time-dependent properties, which condition the delayed response of the cell to forces that are applied and maintained over a longer timescale, are best represented by the cell wall’s viscosity.

A range of methods have been developed to measure the material properties of the cell wall, with indentation techniques providing high spatial resolution and, thus, highly localized information in a non-invasive fashion (3, 12–14). Cellular Force Microscopy (CFM) is a microindentation system specifically developed to measure global mechanical properties of living and growing cells (15). Independent of the type of loading placed on a cell, this technique yields the force measured by the probe on the cell wall as a function of the measured indentation depth. The apparent stiffness of the cell can directly be extracted from the force–indentation curve obtained from short timescale indentation experiments by evaluating the slope of its linear fit (Figure 2). In combination with geometric parameters, such as the cell diameter before and after plasmolysis and the thickness of the cell wall determined by Transmission Electron Microscopy (TEM), mechanical parameters (e.g., elastic modulus and turgor pressure) can be extracted using Finite Element Method (FEM)–based inverse modelling (16). Alternatively, relaxation experiments at longer timescales yield force versus time curves that can be fit to obtain time-dependent material information, such as viscosity (Figure 2). The main challenge, however, lies in closing the loop by relating such mechanical properties to the biochemical composition of the cell wall in order to explain and predict cellular morphogenesis. Here, we achieve this goal by combining short- and long-timescale indentation experiments on PTs of several cell wall mutants in the plant model *Arabidopsis thaliana*. PTs are a well characterized system to investigate cytomechanical properties. Employing the extensive mutant collections in *Arabidopsis* allowed us to correlate variation in the biochemical composition of the cell wall with changes in its mechanical properties and PT morphogenesis.

Our aim was to develop a model that would allow us to distinguish between PTs with seemingly similar phenotypes (growth aberrations and premature bursting) evoked by mutations that affect different aspects of the PT cell wall. Specifically, we investigated the effects of mutations targeting xyloglucan (XyG) and pectin polysaccharides, as well as extensins, a family of cell wall proteins. XyGs are important for the formation of tight associations with cellulose microfibrils, so called ‘biomechanical hot spots’ (17, 18). The regulation of this network alters the dynamics of cell wall elastoplasticity, modulates its extensibility, and allows for controlled cell growth (19–21). The PT cell wall is somewhat special in its composition, as it consists of a very high proportion of pectins compared to the cell wall of other cell types; at the PT tip, it is the predominant cell wall polysaccharide (22, 23). Newly synthesized pectins are methylesterified when they are secreted at the tip. De-esterification mediated by pectin methylesterases (PMEs) and the subsequent crosslinking of acidic pectins by Ca^2+^ reinforce the subapical PT cell wall and play an important role in viscoelasticity (24–27). Taken together, pectins and their esterification state are proposed to be a key determinant in cell wall loosening during cell expansion. Extensins form long rods that make crosslinks with other cell wall components (28). While originally thought to have a largely structural function in the cell wall, it was recently shown that extensins and related proteins play a fundamental role in cell wall integrity signalling (29–32). The differences in cell wall biochemistry between tip and shank alluded to above have been convincingly demonstrated on several occasions (8, 33–36). This led to the widely accepted assumption that the PT tip must be softer than the shank. However, due to the lack of suitable instrumentation, this difference has never been experimentally confirmed. In fact, a softer cell wall at the tip should lead to an expansion of the PT during growth, but wild-type PTs show self-similar growth, maintaining their diameters over time. In earlier work, we could indeed show that the previously reported differences in apparent stiffness between tip and shank (12, 37) can be fully explained by PT geometry and that the assumption of a largely uniform cell wall is compatible with a realistic model of self-similar PT growth (16).

The large variability of biological measurements is often a problem when it comes to modeling. Typically, mean or median values are used as input parameters of the model, such that the model outputs are point estimates of the quantities of interest. As a consequence, the statistical rigor that is applied to the analysis of the experimental data does not carry over to the mathematical models. Because of the often large variability within samples, rigorous statistical methods must also be applied to the models to corroborate the predicted differences between samples. Therefore, a major focus of this work is the reconciliation of the statistical precision in the acquisition of experimental data with uncertainty quantifications of the model.

We employed CFM in combination with FEM-based modelling and a Monte Carlo uncertainty quantification approach to assess turgor pressure and cell wall elasticity in growing mutant PTs. Depending on the nature of the biochemical changes, these parameters were affected differently. Interested in how these changes translate to differences in PT growth, we developed a method to predict the growth rate of PTs based on their cellular mechanical parameters. Since PT growth is driven by time-varying properties, predicting the growth rate requires measuring the viscoelastic properties of the cell wall. Using our combined approach, we were able to show that changes in cell wall biochemistry directly affect growth-controlling parameters, such as turgor pressure and cell wall elasticity. This has an immediate impact on the growth rate of mutant PTs, which we were able to accurately predict and experimentally confirm. Thus, our systems biological approach allowed us to link the biochemical composition of the cell wall to the PT’s growth behavior, which is central to successful plant reproduction.

## Results

### Pollen Tubes of Cell Wall Mutants Have Severe Growth Defects

Under the assumption that mutations in major cell wall components will upset the balance between turgor pressure and cell wall stiffness leading to destabilized PT growth, we analyzed the PT growth phenotypes of different *Arabidopsis* mutant lines covering different components of the PT cell wall. The first line we looked at is deficient in XyG and carries mutant alleles of both *XYLOGLUCAN 6-XYLOSYLTRANSFERASE1* (*XXT1*) and *XXT2* (38). We also analyzed the *xyloglucanase113* (*xeg113*) mutant, which interferes with the arabinosylation of extensins; a functionally relevant feature of these proteins (29). Extensins potentially play a stabilizing role in the absence of XyG (39). Given this possible functional relationship, we also analyzed PTs of the *xxt1 xxt2 xeg113* triple mutant. Since neutral domains of pectins have been shown to bind cellulose microfibrils *in vitro* (40) and, thus, could have a wall stabilizing function, particularly in the absence of XyG (18), we also included the *pectin methylesterase48* (*pme48*) mutant in our studies. The *pme48* mutant affects the de-methylesterification of pectic homogalacturonans (HGs) and, hence, the rigidification of the cell wall through Ca^2+^-crosslinking of HGs (41). A previous study reported high frequencies of morphological abnormalities in *pme48* PTs, including bifurcated PTs with additional tip-growing foci and bursting PT tips (42), reminiscent of the phenotypes we found in *xxt1 xxt2* and *xeg113* PTs (Figure 1).

**Fig. 1.**
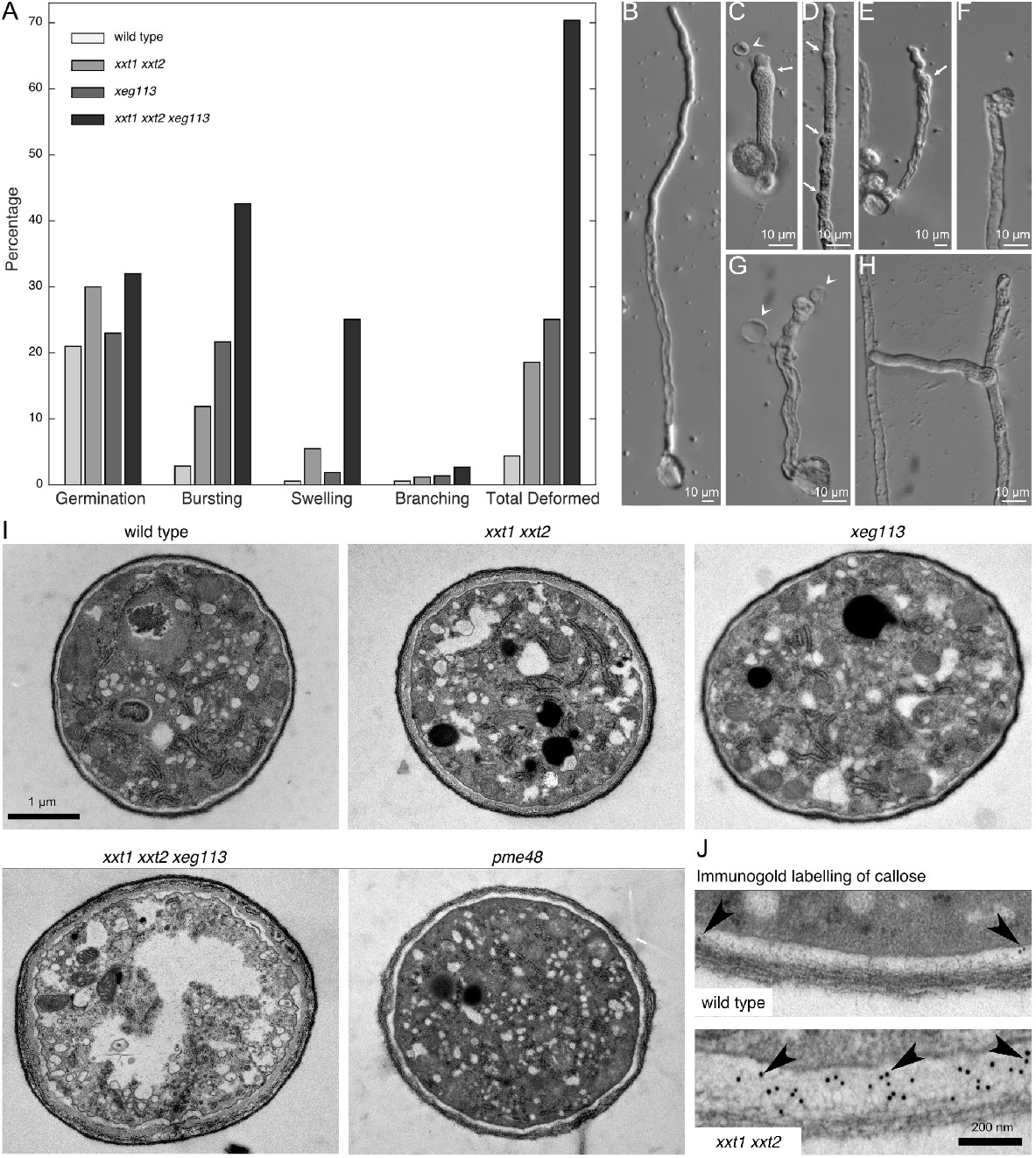
Phenotypes of wild-type and mutant pollen tubes. (A) Quantitative representation of pollen germination and the phenotypes depicted in (B-H).The *pme48* mutant is not shown because the experiment was performed separately and cannot be directly compared. For details see Table S1. (B) Typical wild-type *Arabidopsis* PT. (C-H) Aberrant phenotypes in mutant PTs, such as swelling (C-E), bursting (F), releasing cytoplasmic content (C and G, arrowheads), or branching (H). Tip swelling was often followed by the formation of a new tip, which grew more or less normally (arrows). (C and F) *xxt1 xxt2*, (D) *pme48*, (E and H) *xeg113*, (G) *xxt1 xxt2 xeg113*. (I and J) Cell wall structure of wild-type and mutant PTs. (I) Ultrastructure of transverse sections of wild-type and mutant PTs displaying an outer (dark) and an inner (lighter) layer. Cell wall thickness varies in wild-type and mutant PTs (see also Table 1). The inner wall layer in the *xxt1 xxt2* and *xxt1 xxt2 xeg113* mutants is thickened. In addition, the triple mutant contains clusters of electron-dense material sandwiched between the two cell wall layers. (J) Immunogold labelling (arrowheads point to gold particles) showing a higher accumulation of callose in the inner cell wall of *xxt1 xxt2* PTs compared to the wild type. Scale bars: 10 µm in (B-H), 1 µm in (I), 200 nm in (J).

An examination of the phenotypes caused by changes in cell wall composition revealed a higher germination rate in pollen lacking XyG (*xxt1 xxt2* and *xxt1 xxt2 xeg113*) compared to the wild type (for details, see Table S1). We also observed a significantly higher proportion of burst PTs in these mutant lines (Figure 1A). Among the PTs that were still growing, we found that mutant PTs, especially *xxt1 xxt2 xeg113*, exhibited irregular growth patterns, such as swelling or branching, more frequently than the wild type, which typically formed long, regularly shaped PTs when germinated in liquid pollen tube growth medium (PTGM; Figures 1A to 1H). The XyG-deficient *xxt1 xxt2* PTs showed a stronger tendency for tip swelling than the extensin mutant *xeg113*. The highest frequency of defects, however, was found in the triple mutant *xxt1 xxt2 xeg113*, in which both XyG levels and extensin function are impaired, and 25 % of the PTs had swollen tips (Figure 1A; Table S1). Tip swelling was not always fatal for the PTs because, often, a new tip formed on the club-like, swollen bulge that replaced the original tip, resulting in a temporary resumption of normal growth (Figures 1C to 1E). Especially in *pme48* PTs, tip swelling occurred periodically, leading to local thickenings in otherwise normal-looking PTs (Figure 1D). Branching was relatively rare but, again, occurred at higher frequency in *xxt1 xxt2 xeg113* than in the other mutants (Figures 1A and 1H). The observed irregular growth phenotypes indicate a disturbed balance between turgor pressure and cell wall stiffness, resulting in a strong tendency to burst (Figures 1A and 1F). This is further supported by the fact that the highest bursting rate of 43 % was observed in the mutant disrupting multiple cell wall components, *xxt1 xxt2 xeg113*.

**Table 1.**
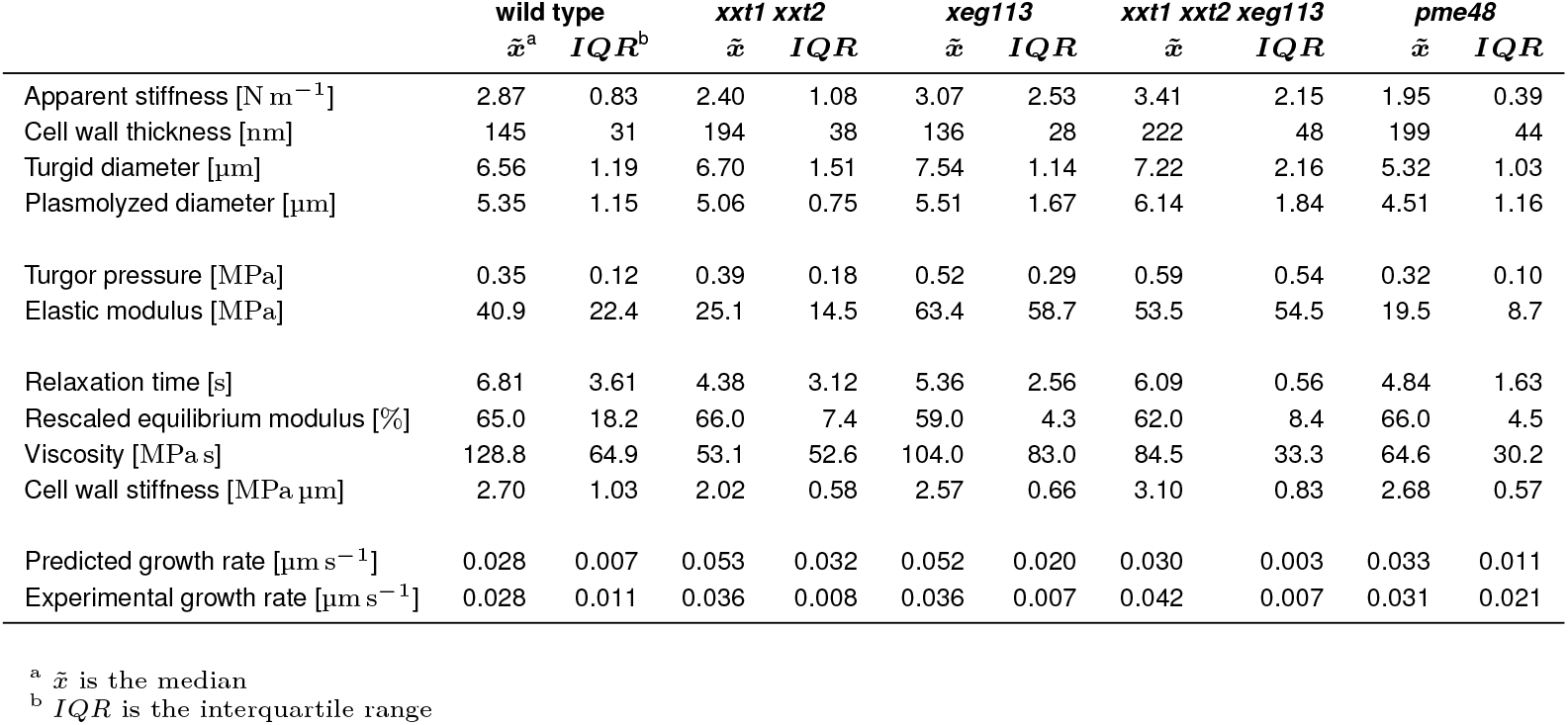
Model input and output parameters.

To determine how the different mutations altered the ultrastructure of the PT cell wall, we used TEM to examine cross sections of PTs grown in a semi-*in vivo* system. The cell wall of wild-type PTs consisted of a dark, fibrous outer layer and a lighter inner layer (43). A similar architecture was observed in the mutant PTs; however, in *pme48* mutants, the outer layer was less dark and the fibrous material seemed more loosely packed, whereas in *xxt1 xxt2* and *xxt1 xxt2 xeg113* mutants, the inner layer was significantly thicker than in the wild type (Figure 1I). The inner cell walls of *xxt1 xxt2* and *xxt1 xxt2 xeg113* PTs are reinforced by increased callose deposition, as revealed by immunogold labelling of TEM sections (Figure 1J), corroborating aniline blue staining results in *xxt1 xxt2* mutants (Figure S1). Furthermore, in *xxt1 xxt2 xeg113* PTs, sandwiched between the inner and outer cell wall was a region of varying electron density, with occasional clusters of electron-dense materials, indicative of major defects in both the composition and architecture of the cell wall (Figure 1I). Given the observed differences in callose accumulation, the susceptibility of the various mutant PTs to callose degradation was investigated. Increasing concentrations of the callose-degrading enzyme lyticase (44) resulted in reduced PT growth, with a clearly stronger sensitivity of *xxt1 xxt2* and *xxt1 xxt2 xeg113* PTs (Figure S2). This result demonstrates the relevance of callose for *Arabidopsis* PT growth, where it can compensate for the absence of XyG in the cell wall of mutant PTs.

### Quantification of Mechanical Properties in Wild-Type and Mutant Pollen Tubes

To elucidate the relationship between the biochemical composition of the cell wall and the mechanical properties that regulate variable growth behavior in the mutant lines studied, several input parameters for the subsequent numerical simulations were measured experimentally (Figure 2A). These parameters include apparent stiffness, cell wall thickness of the turgid PT, and the diameters of turgid and plasmolyzed PTs (Figure 3A and 3B; Table 1).

**Fig. 2.**
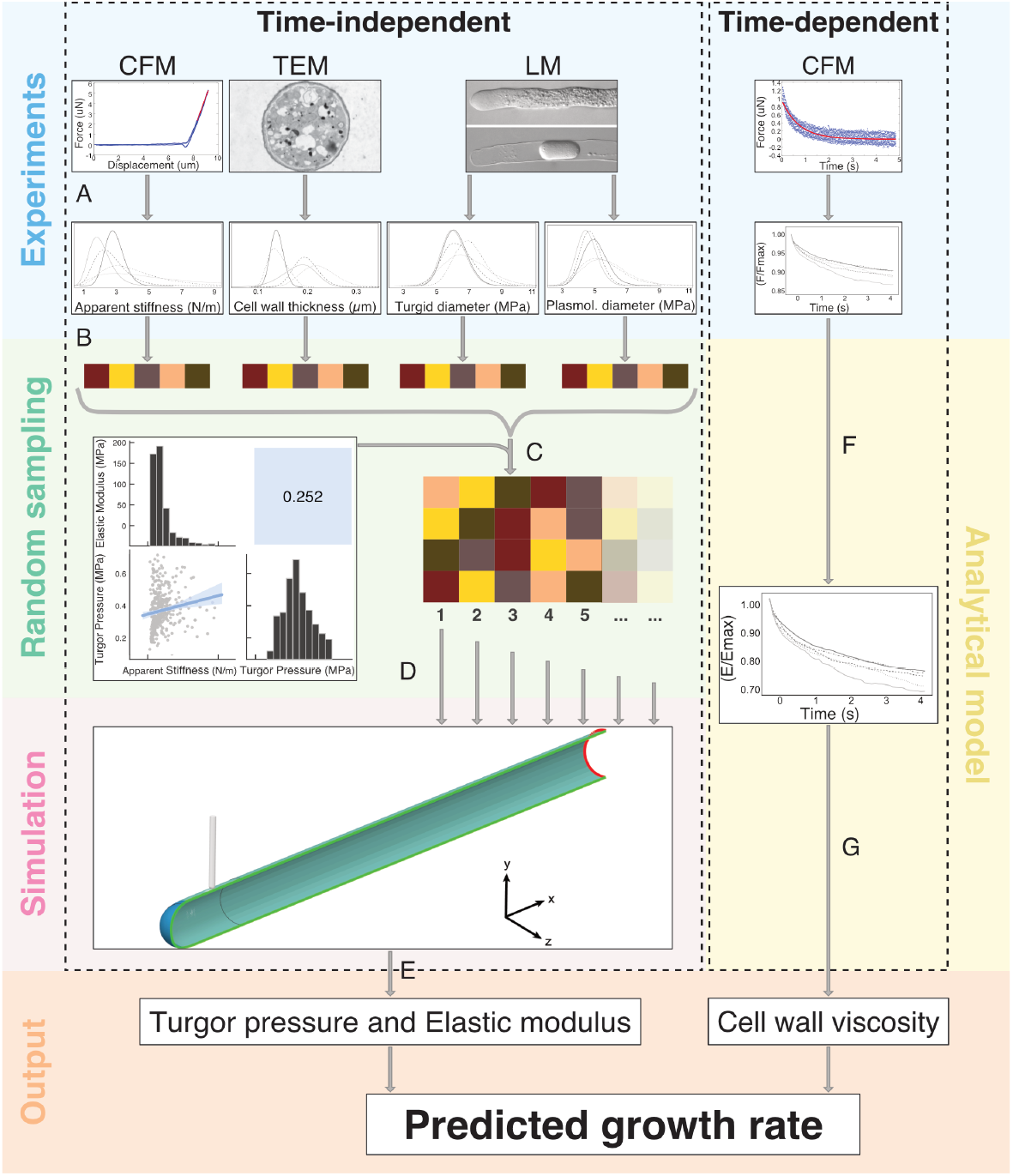
Methodological workflow to predict the pollen tube growth rate. (A, left) CFM, TEM, and light microscopy (LM) experiments to measure the apparent stiffness, cell wall thickness, and the diameter of turgid/plasmolyzed PTs for each variant (wild type and mutants), to which univariate probability distributions are fit. (Right) Stress relaxation experiments using the CFM are performed to extract the normalized force decay in the PT variants over 5 s. (B) Using Latin hypercube sampling, 1000 samples are drawn from the apparent stiffness, cell wall thickness, and turgid/plasmolyzed PT diameter probability distributions for each PT variant (45). (C) A correlation matrix between the apparent stiffness, cell wall thickness, and turgid/plasmolyzed PT diameters is created using inverse FEM modelling of random conformations of the indentation process. This correlation matrix is then used to correlate the samples drawn from the parameter probability distributions to find correlated sets of parameters th are physically possible in the PT system (46). (D) For uncertainty quantification, the correlated sets of parameters are provided as inputs to the FEM-based model where at least 150 correlated parameter sets are simulated per PT variant. (E) The output turgor pressure and elastic modulus distributions for each PT variant are used as inputs in the growth equation (Eq. 3) to predict the PT growth rate. (F) The normalized relaxation modulus decay is estimated from the force decay measurements for all PT variants using equation Eq. S1. (G) Fitting the relaxation modulus decay measurements with equation Eq. S2 yields the cell wall viscosity of the PT variants, which are coupled with the time-independent parameters as inputs in the growth equation.

**Fig. 3.**
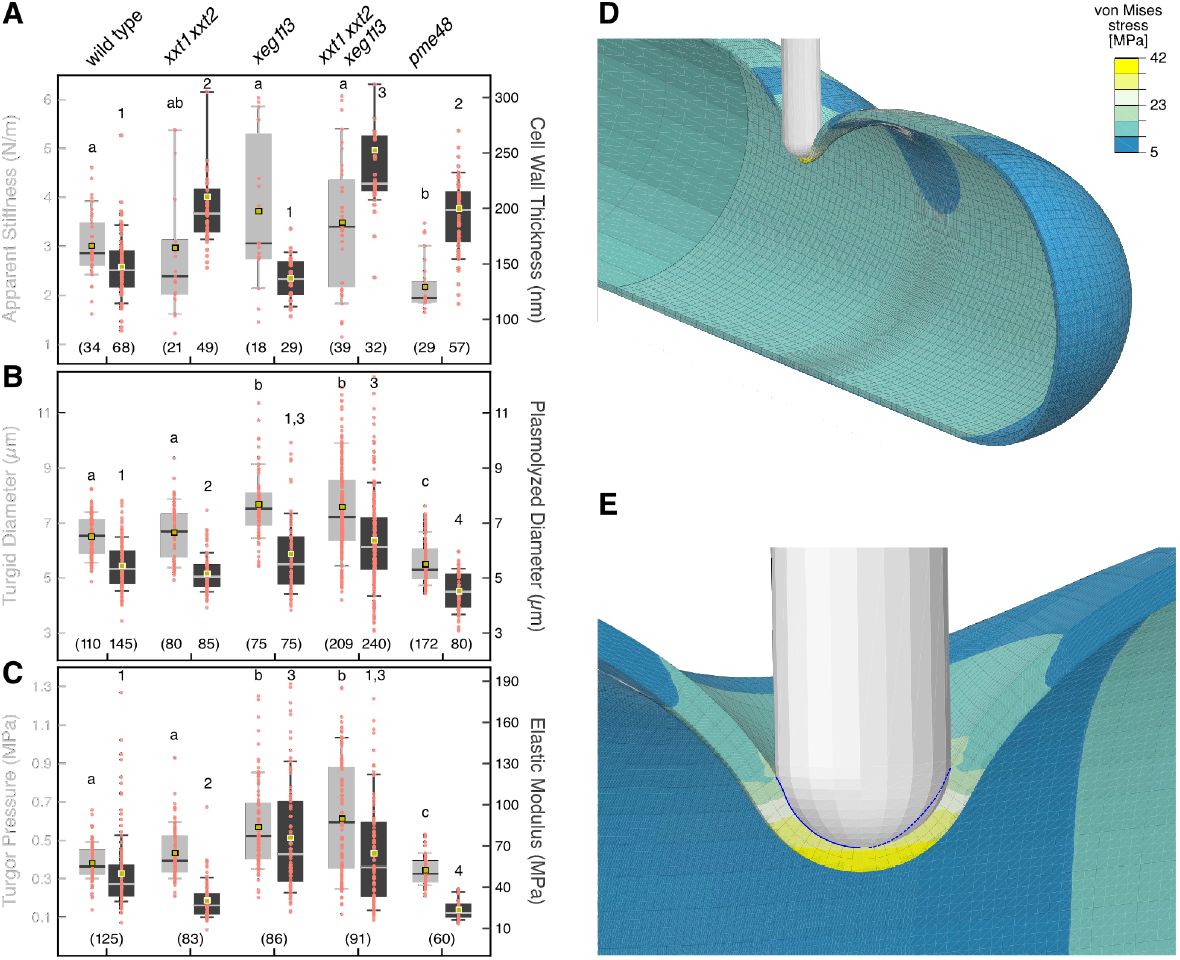
Simulated mechanical properties of wild-type and mutant pollen tubes. (A) Boxplot of apparent stiffness and cell wall thickness of wild-type and mutant PTs measured by the CFM and TEM, respectively. (B) Boxplot of diameters in plasmolyzed and turgid PTs measured with the LM. (C) Boxplot of the output turgor pressure and elastic modulus of wild-type and mutant PTs from the FEM/Monte Carlo simulations. The whiskers in A-C represent the data between the 10^th^ and 90^th^ percentiles. The number of analyzed samples is provided at the bottom of each plot, while significance indicators are located at the top. Medians not sharing a common letter (or number) are significantly different from each other. (D) Finite element mesh of the PT, consisting of a hollow tube constrained to a hemispherical tip, in contact with the rigid indenter of the CFM. A half model is used for computational efficiency. Colors indicate the distribution of von Mises stress caused by the indentation. Mesh density is highest in the indented section to resolve the contact problem and get a high-resolution stress distribution. (E) Close-up of the indented area. The von Mises stress is highest directly underneath the indenter and reduces drastically with distance from the indenter tip. The contact boundary between the indenter and the PT cell wall is indicated by a blue line (dashed where the indenter masks the contact area).

To measure the apparent stiffness, we used the CFM to indent the PTs 10 µm behind the tip in the cylindric part of the subapical region. When using one-dimensional sensors, accurate measurements can only be made perpendicular to the surface of the cell, preventing indentations directly at the spherical tip (3, 16). However, it is legitimate to assume uniform mechanical cell wall properties for the FEM-based model, which we previously validated (16). The CFM experiments revealed that the apparent stiffness was largely variable for all mutant lines except *pme48*, which also displayed significantly lower values (Figure 3A). Cell wall thickness, a critical parameter that determines how much tensional stress can be withstood at a given stiffness, was revealed to be similar in wild-type and *xeg113* PTs. All other mutants, however, had values that significantly exceeded those of the wild type, whilst diverging significantly from each other. Since the cell wall of a turgid PT is constantly under tension due to its internal turgor pressure, PT plasmolysis leads to considerable radial shrinkage. The magnitude of this shrinkage acts as a proxy to characterize the elastic behavior of the PT cell wall (16). Wild-type and *xxt1 xxt2* PTs displayed similar turgid diameters, while *xeg113* and *xxt1 xxt2 xeg113* mutant PTs exhibited significantly larger and *pme48* PTs significantly smaller diameters, respectively (Figure 3B). For plasmolyzed diameters, only *xxt1 xxt2 xeg113* and *pme48* mutant PTs showed significantly higher and lower values than the wild type, respectively. These measured parameters were sequentially fit to yield the elastic modulus and turgor pressure of the PTs (Figures 3C to 3E), using an inverse continuum FEM-based model of CFM indentation (47).

### Modelling Mechanical Input Parameters

Due to large intra-sample variance and skew in the experimentally determined input parameters, an elaborate Monte Carlo uncertainty quantification was performed within the FEM framework, starting with a parametric fit on all experimental observations (Fig. 2B). The fitted probability distributions were then sampled with Latin hypercube sampling (45), and the resulting samples were correlated using the Iman–Conover method (46, Figure 2C). Input parameter realizations from these correlated distributions were subsequently simulated until convergence (minimum 100 realizations) for each mutant using the inverse FEM-based model, until the averaged PT elastic properties—turgor pressure and elastic modulus—could be extracted (Figures 2D to 2E).

Since a large number of input realizations had to be simulated to obtain accurate statistics for turgor pressure and elastic modulus, the FEM-based model was set up to ensure robustness and computational efficiency, whilst not sacrificing on accuracy (for a detailed description of the model see supplemental data).

The model output showed that the *xeg113* and *xxt1 xxt2 xeg113* turgor pressure values were significantly elevated, while the turgor pressure in the *pme48* mutant was lower than in wild-type PTs. Furthermore, the elastic moduli of *xxt1 xxt2* and *pme48* PTs were significantly lower compared to the wild type, while those of *xeg113* and *xxt1 xxt2 xeg113* PTs were significantly higher. The turgor pressure and elastic modulus interquartile ranges (IQR, *x*25−*x*75) for *xeg113* and even more so for *xxt1 xxt2 xeg113* PTs doubled compared to the wild type (Figure 3C), indicating a disruption of the fine-tuned regulation of cellular morphogenesis. Similarly, the lowered apparent stiffness, turgor pressure, and elastic modulus of *pme48* PTs fit well with the predicted effect of a less rigid cell wall resulting from reduced HG de-esterification.

### Principal Component Analysis (PCA) Indicates That Apparent Stiffness, Stretch Ratio, and Cell Wall Thickness Are Key Parameters of Pollen Tube Growth

As the correlations between input and output parameters were *a priori* unknown, random realizations of all parameters in the IQR (*x*25−*x*75) were simulated to obtain a correlogram (Figure 4A). Clearly, the apparent stiffness was more correlated to turgor pressure than any other parameter, indicating that the CFM technique is most sensitive to fluctuations in turgor pressure at a calibrated indentation depth and osmolarity of the medium. The elastic modulus was shown to correlate positively with the turgid diameter and negatively with the circumferential stretch ratio (the ratio between turgid and plasmolyzed diameter). Furthermore, cell wall thickness and elastic modulus were negatively correlated, indicating a synergistic relationship that is consistent with cell wall remodeling under external stress, corroborating results from other biophysical analyses (48).

**Fig. 4.**
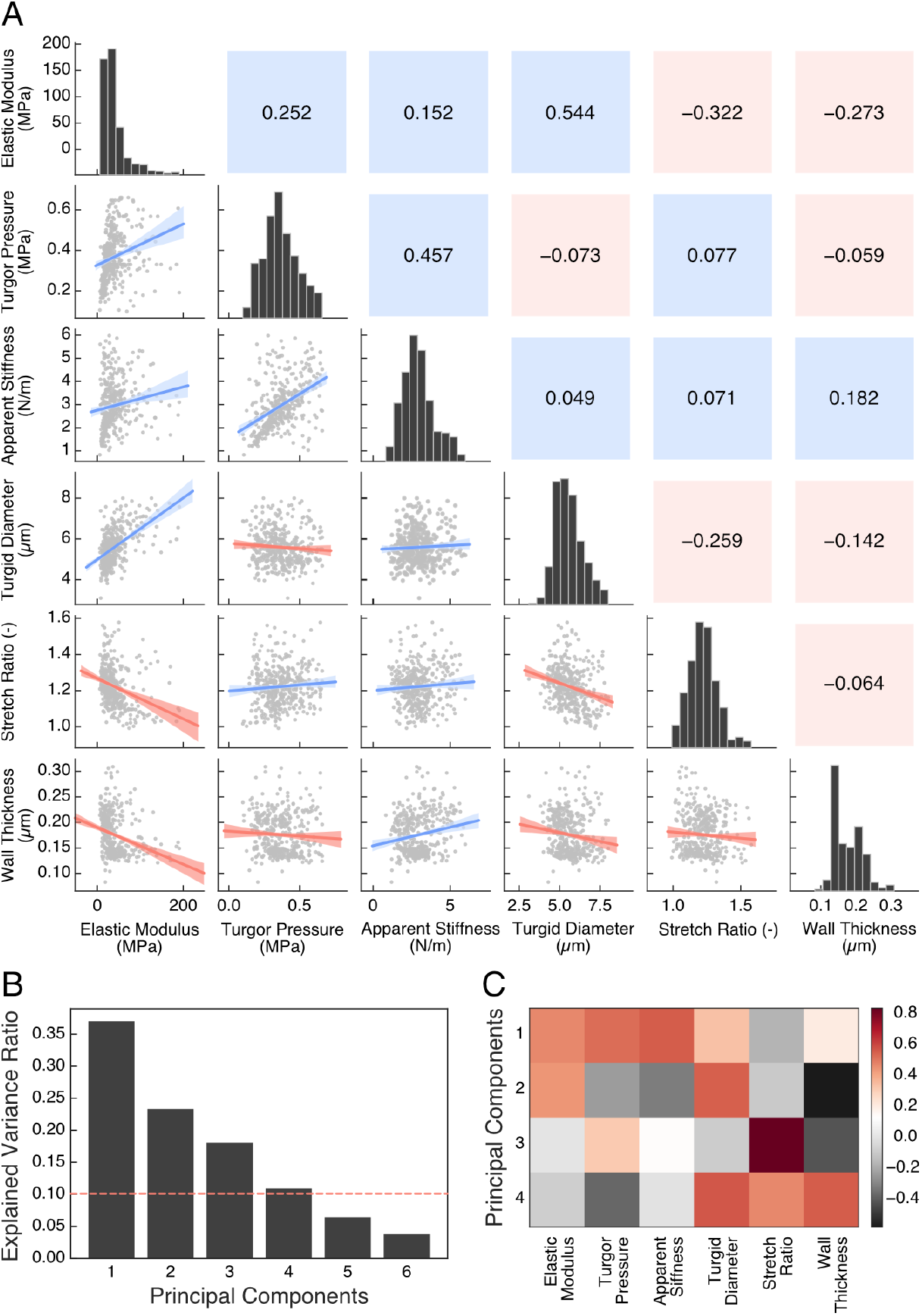
Correlation and sensitivity analysis of the input and output parameters of the FEM/Monte Carlo simulations. (A) Correlogram of the parameter space including Pearson’s correlation coefficients (upper triangle), histograms (diagonal), and correlation scatter plots (lower triangle) between parameters. Blue indicates a positive and red a negative correlation, respectively. (B) Explained variance ratio of the input/output parameters based on PCA. Three components are responsible for ∼80 % of the variance in the system. (C) Contribution of input/output parameters to PCA components. Apparent stiffness, circumferential stretch ratio, and cell wall thickness measurements provide most information gain.

To study the degree of variability in the parameter space, Principal Component Analysis (PCA) was performed on the input and output variables (Figures 4B and 4C). Unexpectedly, only three components accounted for ∼80 % of the variability in the system, with apparent stiffness, circumferential stretch ratio, and cell wall thickness measurements providing most information gain. This indicates that the relatively simple shape of the PT does not introduce complex inter-dependencies between the input and output parameters. Furthermore, while the elastic modulus is clearly crucial to the regulation of PT growth, this analysis shows that, under our model assumptions, its variability is largely explained by other measured parameters. While the cause-and-effect relationship between these parameters is yet unclear, direct experimental measurements of the elastic modulus seem to be unnecessary to accurately determine PT growth rates.

### Measuring the Viscosity of the Cell Wall

The viscosity of the cell wall plays a central role in regulating the PT growth rate (49–53). Biophysical methods, such as the microindentation of static tissue to induce stress-relaxation, have previously been used to quantify viscosity (13, 54, 55). Stress-relaxation is defined as a time-dependent reduction in stress when a material is subjected to a constant mechanical strain, usually caused by rearrangements in the conformation or position of load-bearing polymer chains. This process occurs on multiple timescales corresponding to the degree and type of polymer chain displacement. The difference in viscosity between cell wall mutants is probably caused by modifying essential components of the cell wall, like pectins and XyG, that have been shown to impact time-dependent properties (56). Therefore, an accurate estimation of the relative growth rate between *Arabidopsis* cell wall mutants is likely to strongly depend on cell wall viscosity, which thus needs to be accurately quantified in wild-type and mutant PTs.

For the purposes of this analysis, PTs were indented with the force sensor of the CFM until a maximal force of 5 µN was reached and held at this position while recording the force decay (Figure 5A). As the PTs were growing throughout this indentation procedure, the experiment faced two notable constraints compared to previous studies (54, 55). Firstly, since the growing PT is a dynamic system that absorbs water from the surrounding medium through osmosis (57), we limited the applied strain duration to 5 s in order to avoid secondary effects that could influence the force measurements (Figure 5B). For PTs of all lines, the measured force was found to decay to a constant level in less than 5 s, indicating that the applied strain duration was sufficient to capture the time-dependent properties relevant for PT growth (Figure 5C). To rule out measurement artifacts, a purely elastic substrate and a highly viscous glass-fiber were indented as controls using the same protocol. As expected, the force decay curves of the PTs lay within those two extremes. Secondly, as in short timescale CFM analyses, the measured force must be converted to stress to extract the time-dependent equivalent of the elastic modulus, known as the relaxation modulus. Using a simulation approach to model this experiment is not straightforward because a continuous forcing function like turgor pressure coupled with a viscoelastic cell wall renders FEM modelling unfeasible without including material plasticity to limit radial strain. However, measuring cell wall plasticity in a living and growing PT is simply impractical with currently available methods. To circumvent this constraint in estimating the relaxation modulus, we used an analytical relation developed for the nano-indentation of thin shell tubes in combination with the turgor pressure obtained from the FEM simulations.

**Fig. 5.**
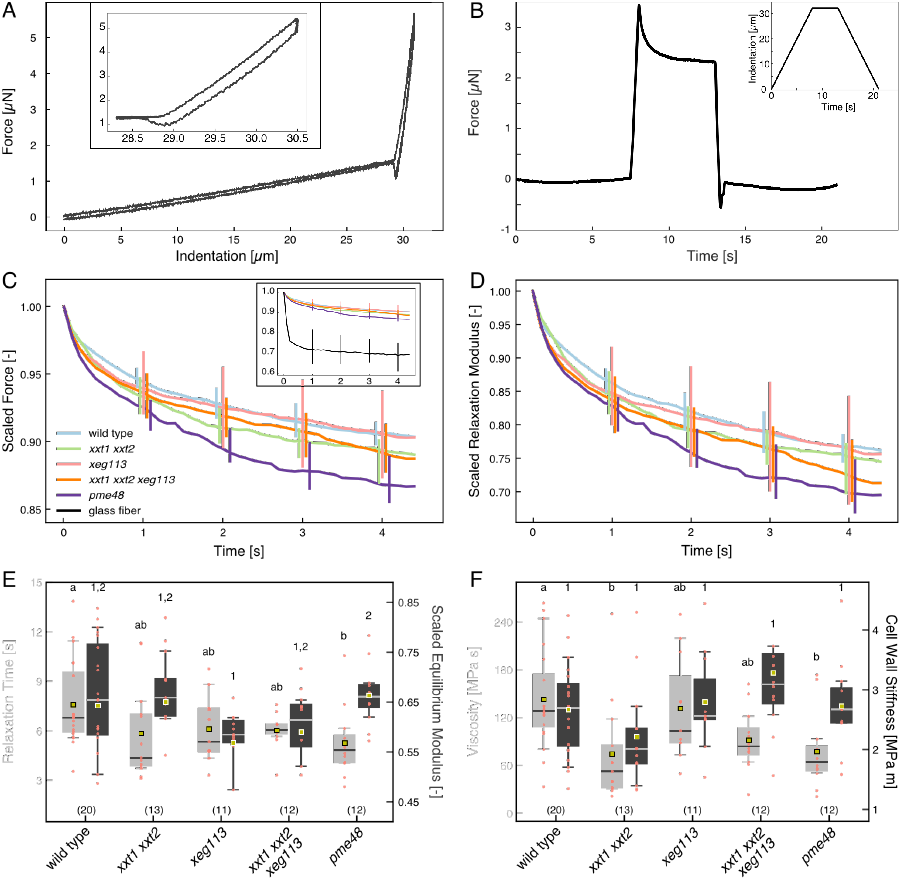
Experimental indentation results to measure cell wall viscosity. (A) Force-Indentation and (B) Force-Time output from CFM, illustrating 5 s stress-relaxation on the PT by the CFM force sensor. The inset in (A) shows a closeup of the curve where the sensor is in contact with the PT. Note that the curve in the inset is corrected for the effect of the force increase while the sensor is approaching the PT. The inset in (B) shows the vertical movement of the force sensor during the experiment. (C) Force decay (scaled by the initial force at *t* = 0) during the retraction phase of the stress relaxation experiment on wild-type and mutant PTs and a control substrate (glass-fiber, shown in the inset). (D) The relaxation moduli (scaled by the initial relaxation moduli at *t* = 0) of wild-type and mutant PTs found with equation Eq. S1, exhibiting a two-phase relaxation process. (E) Comparison of relaxation times *τ* and equilibrium moduli (scaled by the initial relaxation moduli at *t* = 0) from the fitting equation Eq. S2. (F) The viscosity derived from *τ* = *η/E*? compared with the wall stiffness obtained from Figure 3 (E). The error bars in (C) and (D) were spread apart along the time axis for better distinguishability. Originally, they would all overlap at the position of the *xeg113* error bar. The whiskers in (E) and (F) represent the data between the 10^th^ and 90^th^ percentiles. The number of analyzed samples is provided at the bottom of each graph, while groups indicating significance are located at the top.

As seen in Figure 5D, the relaxation modulus of the cell wall relaxes in two distinct phases; the initial fast decay (*t<*0.5 s) followed by the long-term slow relaxation (*t>*0.5 s), consistent with previous studies on plant tissues (13, 55). This observation necessitates the use of two superimposed relaxation times *τi*. However, since the slow relaxation time corresponds to the timescale of PT growth, this value, along with the equilibrium modulus, is the main parameter for PT growth (Figures 2G and 5E). From these parameters, we can estimate the viscosity of the cell wall for wild-type and mutant PTs and weigh it against the stiffness of the cell wall (*Eh, h* being cell wall thickness) found by the elastic FEM models (Figure 5F).

The relaxation times extracted from the stress-relaxation experiments showed a large statistical dispersion between 2 and 14 s even though the median values for the different mutants were constrained within a relatively small interval of around 5 s (Figure 5E). Some outliers are expected because, due to the inhomogeneity of the cell wall surface, the indenter is likely to occasionally indent local wall structures which are more pliable than the bulk cell wall material, resulting in outliers with higher relaxation times. Overall, the wild type displays the highest relaxation times, with only the *pme48* consistently showing significantly lower relaxation times. The equilibrium modulus is found to be consistent between wild-type, *xxt1 xxt2*, and *pme48* PTs at 65 %, compared to *xeg113* and *xxt1 xxt2 xeg113*, which returned slightly lower values. These findings indicate that the proportion of elastic-like behavior in mutant PTs are not significantly affected by their altered cell wall composition. *xeg113* PTs exhibit a distinctly shaped relaxation modulus decay curve that relaxes similar to the other mutants in the initial fast decay but returns to a wild-type-like behavior in the long-term slow relaxation phase (Figure 5D). This similarity between wild-type and *xeg113* PTs is also reflected in the equivalence of their viscosities, which might indicate that, if any fluid-like structural changes are caused by the *xeg113* mutation, they are adequately compensated for by other cell wall components (Figure 5F). *xxt1 xxt2* and *pme48* mutants show similarly large drops in viscosity compared to the wild type, pointing to a significant increase in the fluid nature of their cell wall. The viscosity of *xxt1 xxt2 xeg113* lies between *xxt1 xxt2* and *xeg113*, which may indicate a unique cell wall structure with only partial compensation for the loss of XyGs. Indeed, a lack of XyGs appears to also strongly affect the elastic properties of the cell wall in *xxt1 xxt2* PTs, which display a significantly lower wall stiffness than the wild type and other mutants, despite *xxt1 xxt2* PTs possessing a thick cell wall without obvious structural defects.

### Mechanical Tip Growth Models Accurately Predict Growth Rates of Wild-Type and Mutant Pollen Tubes

Tip growth has been studied for decades by building models predicated on assumptions with varying degrees of complexity. These models include the pre-eminent Lockhart model, based on the assumption of a viscoplastic cell wall (49), an extended version accounting for calcium dynamics, variations in cell wall thickness, and material extensibility at the tip (58), a further extension including the anisotropy of the cell wall (50), simulating spatially confined growth (4), and a model focusing on stress-relaxation due to loss of stability (59, 60). The consensus among these models is that the cell wall contains both elastic and viscous components and can, therefore, be described by a linear viscoelastic constitutive law. The Lockhart model in particular utilizes the concept of a Bingham plastic fluid to describe cell wall mechanical properties. These are non-Newtonian fluids that behave like rigid bodies at low stress but flow like a viscous liquid at high stress, such as mud or mayonnaise. Applied to the cell wall, it manifests as a material that flows like a fluid (or grows uniaxially in the case of the PT) when subjected to turgor pressure above a given yield pressure but sustains no permanent deformation at lower turgor pressures. The contribution of the elastic deformation of the cell wall to growth was later augmented to the Lockhart model (49) by Ortega and colleagues (61). Assuming isotropic material properties, the strain rate of the cell wall 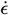 is described by

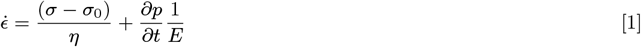

where *σ* is the wall stress, *σ*0 is the yield stress, *η* is the viscosity, and *E* is the elastic modulus. Previous studies using pressure probes have indicated that the turgor pressure in *Lilium longiflorum* PTs is stationary over time, with small pressure fluctuations proving to be uncorrelated with the growth rate (9, 62). For this analysis, we therefore assume that *∂p/∂t* = 0. To extend the strain rate relation into the volumetric growth rate *v* of the PT, the PT tip is parametrized as a hemispherical shell with diameter *d* as its characteristic length scale and wall stress *σ* given by

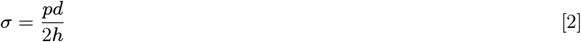

at turgor pressure *p* and cell wall thickness *h*. The average volumetric growth rate *v* of the PT can thus be described by

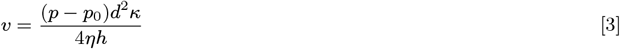

where *v* is the product of the strain rate and the radius (*d/*2), *p*0 is the yield pressure, and *κ* is a scaling factor. Lockhart’s growth equation only captures the equilibrium growth rate, while disregarding the osmolarity of the growth medium, which has been shown to be a strong growth rate regulator (63). Since we aim to compare the relative growth rates between cell wall mutants of *Arabidopsis* under identical conditions, this equilibrium growth rate is sufficient for our analysis. To allow for an equal comparison of the predicted growth rate with the experimental growth rate, we introduced the scale factor *κ* to correct for the effect of the growth medium osmolarity. *κ* is calculated as the ratio of the medians of predicted and experimental growth rates in the wild type and, therefore, simply normalizes the predicted and experimental growth rates for all mutant PTs. A further consideration is that PT growth cycles are known to be driven by the oscillatory nature of viscosity and cell wall thickness (51). However, since we are interested in the equilibrium growth rate over longer timescales, only the average viscosity and cell wall thickness measured by the CFM and TEM experiments are required. Additionally, for the purpose of this analysis, the yield pressure is assumed to be constant across all cell wall mutants (51). Nevertheless, it should be noted that the yield pressure of *pme48* mutant PTs is more likely to diverge from the wild type compared to the other mutants because *pme48* impairs the crosslinking of pectin, the major cell wall component at the PT tip (5).

To challenge the model prediction, we determined the growth rate experimentally by tracking PT growth for 20 min and averaging it over time (Figures 6A and 6B). The experimental growth rate of *xxt1 xxt2* PTs was significantly higher than that of the wild type (∼50 %), which is also reflected in the predicted growth rate (Figures 6C and 6D). The more fluid-like nature of the *xxt1 xxt2* cell wall coupled with its lower cell wall stiffness provides a clear rationale for the elevated growth rate of this mutant. The similarly high growth rate (∼50 % above the wild type) of *xeg113* was also effectively captured by the growth model. However, since the viscosity of *xeg113* is similar to wild type, this growth rate elevation is more likely resulting from other factors, namely the significantly higher turgor pressure and diameters of *xeg113* PTs. Despite overestimating the increase in PT growth rates compared to the wild type, the growth model predictions for *xxt1 xxt2* and *xeg113* mutant PTs were found to be similar to each other, mirroring their experimental growth rates. This indicates that the model accurately weighs the opposing actions of viscosity, turgor pressure, and cell wall thickness on the PT growth rate. *xxt1 xxt2 xeg113* PTs display the fastest experimental growth rate (∼80 % higher than the wild type), seemingly indicating an additive effect of the *xxt1 xxt2* and *xeg113* mutations. The predicted growth rate of *xxt1 xxt2 xeg113*, surprisingly, resembles that of the wild type. While this might indicate a deficiency in the model, it should be noted that *xxt1 xxt2 xeg113* PTs have a lower viscosity than the wild type, combined with the highest turgor pressure of all analyzed mutants. Therefore, the explanation for its model-predicted growth rate is based solely on its exceptionally thick cell wall. As previously stated, TEM analysis showed that *xxt1 xxt2 xeg113* PTs possess a thick inner cell wall with several architectural defects. Since the growth rate equation considers the cell wall to be a continuum of both the outer and inner cell walls, we re-analyzed the CFM experiments for *xxt1 xxt2 xeg113* PTs considering only cell wall areas that do not show the electron-dense defects. The use of this revised cell wall thickness measurement is likely a better reflection of the growth rate of the *xxt1 xxt2 xeg113* mutant, as it only includes the load-bearing constituent of the cell wall. As anticipated, the resulting predicted growth rate is much closer to the experimental growth rate (Figure 6D). The experimental growth rate of *pme48* PTs displayed a large variability as would be expected when disrupting the main constituent of the cell wall at the PT tip (5). However, the median growth rate was 20 % higher than that of the wild type, which is consistent with previous studies (42). Furthermore, the growth equation successfully balanced the lower cell wall viscosity of *pme48* with the lower turgor pressure to produce an accurate prediction, with no complications arising from the yield pressure assumption.

**Fig. 6.**
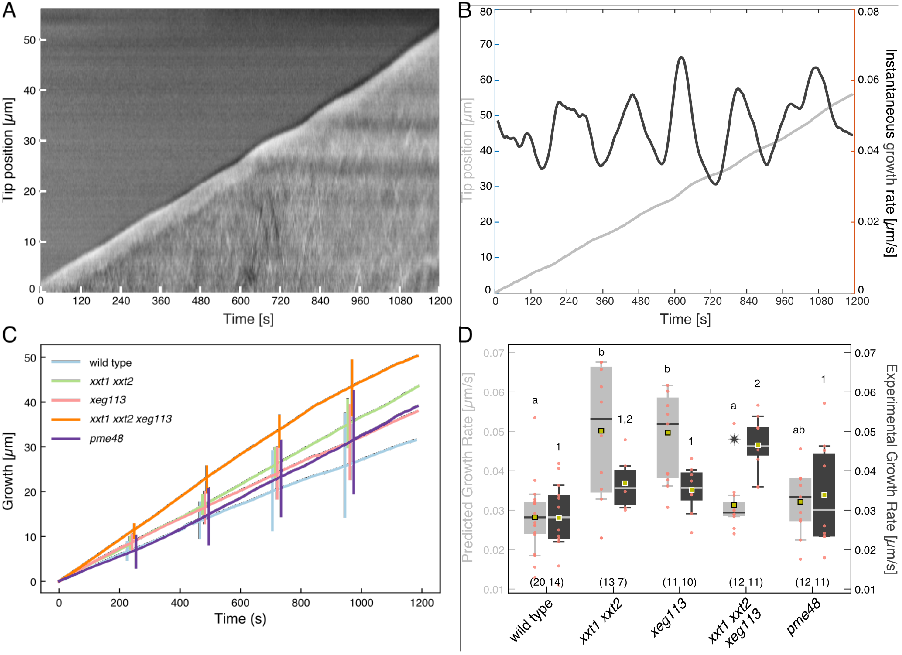
Growth rate analysis of wild-type and mutant pollen tubes. (A) Kymograph of an *xxt1 xxt2 xeg113* PT. (B) Tip position as a function of time and instantaneous growth rate of the PT in (A). (C) Average measured growth of the cell wall mutants over 1200 s. (D) Comparison of simulated and measured growth rates of wild-type and mutant PTs. For *xxt1 xxt2 xeg113* PTs the real growth rate is substantially underestimated by the simulation. Therefore, the star marks the median predicted growth rate when a cell wall thickness of 150 nm for *xxt1 xxt2 xeg113* PTs is assumed. See subsection for discussion. The error bars in (C) were spread apart for better distinguishability. Originally, they would all overlap at the position of the *xeg113* error bar. The whiskers of the boxplots represent the data between the 10^th^ and 90^th^ percentiles. The number of analyzed samples is provided at the bottom, while groups indicating significance are located at the top.

## Discussion

The plant cell wall plays a major role in regulating cell expansion. Loosening the links between the cell wall components just the right amount allows turgor pressure to increase the cell’s volume, whereas timely reinforcement of the thereby weakened cell wall matrix is necessary to avoid bursting. An imbalance between these processes either prevents cell expansion if cell wall loosening is insufficient, or leads to the bursting of the cell if the stabilisation after expansion is not adequate. Thus, a tight coordination between the external cell wall and physiological processes inside the cell is required, particularly in the vacuole, which plays a crucial role in controlling turgor pressure (64). Therefore, a premature bursting of PTs can either be attributed to fluctuations in turgor pressure or to altered cell wall properties (63, 65). In order to investigate how changes in the composition of the cell wall affect its mechanical properties, we developed powerful measurement and simulation tools, which allowed us to discriminate between the similar PT growth phenotypes of several *Arabidopsis* mutants that have a high tendency to rupture. Time-independent indentation experiments in combination with a FEM modelling approach revealed that mutations in primary cell wall components not only affect the elasticity of the cell wall but also influence the magnitude of turgor pressure. While these two parameters were affected differently in the mutants, they generally led to a disturbance of the balance between growth-promoting and growth-limiting factors and, eventually, to the bursting of the PTs. Hence, while the wild type can effectively regulate turgor pressure and cell wall rheology to control PT growth, alterations in turgor pressure or cell wall stiffness in mutants with an altered biochemical composition of the cell wall drastically affect PT growth.

Employing FEM modelling to translate these measured parameters into usable constitutive properties was an essential step in achieving acceptable estimates for the PT growth rate. To build the FEM model presented here, we could benefit from previous efforts that used FEM to model pollen tubes, which, despite some flaws, were seminal for our own approach. An early FEM-based indentation model assumed that the turgor can be completely ignored when indenting a pressurized cell, which contradicts the basic principles of mechanics (66). The first attempts to model PT growth with FEM considered growth to be a purely elastic process (10, 67), resulting in the shank being about 250 times stiffer than the tip to maintain self-similar growth (67), which is highly unrealistic. Taking these shortcomings into account, we changed our previous model (16) and adapted it to *Arabidopsis* PTs (for details, see SI). Biological measurements at the micro-scale are fundamentally subject to significant dispersion. However, in contrast to the situation in wild-type PTs, mutations in cell wall components cause unpredictable growth abnormalities by disrupting the mechanical equilibrium. This imbalance increases until the PT eventually bursts. Uncertainty about the state of mechanical equilibrium at the time of measurement leads to markedly greater intra-mutant variability than can only be reconciled by proper error estimation methods. Therefore, the Monte Carlo simulation-based uncertainty quantification was equally necessary as the FEM model to derive more precise outcomes. Despite using these techniques, the growth rate prediction of *xxt1 xxt2 xeg113* based solely on simulated data was considerably lower than the experimentally determined growth rate. While a closer analysis allowed us to identify a potential source of this disparity, defects that are difficult to directly quantify and capture in the models will always impact the accuracy of such analyses.

It is of note that our choice of mutants does not affect the outcome of the study, since the scope of this manuscript is investigating the question whether PT growth rate can be predicted based on mechanical parameters of the cell wall. Nevertheless, it would be interesting to see how mutants in other components of the PT cell wall integrity machinery may perform in comparison to our newly devised growth rate prediction tool.

As mentioned previously, all mutant PTs had a significantly increased tendency to burst. However, this happened after a growth period of varying length, during which cell wall integrity was maintained despite irregularities, such as temporary stalling, often accompanied by swelling, branching, and leaking cytoplasm (Figure 1). Consequently, although we experimentally analyzed only intact PTs, it was impossible to determine what state of stability a particular PT was in. This can explain the high variability of the apparent stiffness values in mutant PTs with a defective cell wall. Especially for *xxt1 xxt2 xeg113* PTs, which had by far the highest bursting rate, we assume that we have overestimated the apparent stiffness as only the strongest PTs survived and, thus, could be measured.

But why would PTs that completely lack major cell wall components, such as the XyG-free *xxt1 xxt2* double mutant (38) be able to grow at all? We assume that the cell wall integrity pathway is able to sense the deficiency in certain constituents and compensate the defects to some extent by overproducing other compounds (68, 69). Evidence for this hypothesis comes from immunolocalization experiments, showing that in *xxt1 xxt2* PTs displaying a complete lack of XyGs, extensins were strongly overexpressed (Figure S1). Similarly, the inner cell wall of *xxt1 xxt2* and *xxt1 xxt2 xeg113* PTs is thickened by excessive accumulation of callose (Figures 1I, 1J, and S1). Such compensatory mechanisms are further corroborated by the observation that the lack of both XyG and functional extensins leads to synergistic phenotypic effects (Figure 1A).

Numerous studies have focused on extracting structural and hydraulic properties of PTs under specific conditions, while several others have concentrated more on their biochemical characterization. The PT growth rate provides an easily accessible framework to bridge these two perspectives, allowing to shed light onto the cause and effect relationship that exists between them. To our knowledge, this is the first study that has been able to estimate the relative growth rate among PT mutants and, thereby, to gain insights into their unique biochemical alterations and compensatory mechanisms, using solely indentation and geometric measurements.

In summary, we integrated PT growth measurements, CFM to determine mechanical cell wall properties, and TEM analyses of the cell wall to accurately predict the growth behavior of mutants with an altered cell wall composition using FEM simulations with Monte Carlo-based uncertainty quantification of cellular properties. This combination of experimental and modelling approaches provided novel insights into the interplay between biochemical and mechanical factors controlling cellular morphogenesis. Our systems biological study, which applied statistical rigor not only to experimental data but also to modeling aspects, provides a general strategy that can be applied to many other investigations of cell and tissue morphogenesis.

## Materials and Methods

### Plant Material and Growth Conditions

All plants used were *Arabidopsis thaliana* (L.) Heynh of the Columbia (Col-0) accession. *xxt1 xxt2* seeds (38) were obtained from the Nottingham Arabidopsis Stock Centre (NASC). *xeg113–2* (29) and *pme48* (42) were kindly provided by Markus Pauly and Jean-Claude Mollet, respectively. Seeds were sown on half-strength MS media (1/2 MS salt base, 10 % sucrose, 0.05 % MES, 0.8 % Phytoagar, pH>5.7 with KOH), stratified for 2–3 days at 4 °C in the dark, and then moved to long-day conditions (8 h dark at 18 °C, 16 h light at 22 °C, 60 % humidity). When showing two to four true leaves, seedlings were transplanted to soil and grown under long-day conditions in a walk-in growth chamber (8 h dark, 16 h light, 22 °C, 60 % humidity).

### Pollen Tube Culture Conditions

PTs were grown *in vitro* for all experiments except for the TEM analysis to determine cell wall thickness where we used a semi-*in vivo* approach. For *in vitro* pollen tube growth, flowers were collected and incubated for 30 min at 22 °C in a moisture chamber. Liquid pollen tube growth medium (PTGM; 5 mm CaCl_2_, 5 mm KCl, 1.6 mm H_3_BO_3_, 1 mm MgSO_4_, and 10 % (w/v) sucrose, pH 7.5) was prepared as described (70). Pollen grains were brushed onto silane-coated slides and covered with liquid PTGM, germinated, and grown in a moisture chamber, first at 30 °C for 30 min and then at 22 °C for at least 5 h. For germination on solid medium, 1.5 % low melting agarose was added to liquid PTGM. For CFM measurements PTs were germinated on silane-coated slides. To determine the growth rate, PTs were grown in PDMS-microchannels (71) under the same culturing conditions as for normal *in vitro* growth. For TEM analysis, PTs were grown in a semi-*in vivo* system as described in (43). Sepals, petals and stamen were removed from freshly opened Col-0 flowers. The stigmas were pollinated with pollen from the Col-0 and the mutant lines, respectively. Pistils were cut right below the style and incubated on solid PTGM in a moisture chamber and incubated at 22 °C for at 3.5 h.

### Immunocytochemical Analysis of Pollen Tubes

PTs grown for 5 h on silane-coated slides were fixed in PEM buffer (4 % formaldehyde prepared freshly from paraformaldehyde in 1 m NaOH, 50 mm PIPES, 1 mm EGTA, and 5 mm MgSO_4_, pH 6.9). For the enzymatic digest of selected wall components, fixed PTs were rinsed with sodium acetate buffer (pH 5.5) and incubated with a 5 U*/*mL solution of xyloglucan-specific xyloglucanase prepared in the same sodium acetate buffer at 37 °C for 2 h. Enzyme-treated and non-treated fixed samples were rinsed three times with phosphate buffered saline (PBS) for 5 min each, and blocked with 5 % skim milk powder (MP) in the same PBS buffer for 1 h or overnight at 4 °C. Controls included non-digested samples and/or omitting the primary antibody. The PTs were then incubated with a 10-fold dilution of primary monoclonal Antibodies (mAbs) in PBS containing 5 % (w/v) MP for 1 h. Samples were washed in PBS and incubated with a 100-fold dilution of fluorescein isothiocyanate-labeled secondary Ab (Sigma) in PBS/MP for 1 h in darkness. The samples were washed three times in PBS and mounted in a glycerol-based Citifluor AF1 anti-fade solution. Fluorescence was detected on a Leica DM6000 microscope (excitation: 480*/*40 nm, emission: 527*/*30 nm).

### Lyticase Treatment

For lyticase treatment, increasing concentrations (as indicated in Figure S2) of lyticase were added to the PTGM. Differential interference contrast (DIC) on a Leica DM6000 microscope was used for documentation of growing PTs. The length of the PTs was analyzed using Fiji.

### TEM Analysis

A detailed description of the TEM method has been published elsewhere (43). Briefly, PT specimens were fixed in 1.25 % glutaraldehyde in 0.05 % cacodylate buffer, post-fixed in 2 % OsO4, dehydrated in acetone, and embedded in Epon. Then thin sections from between 5 to 15 µm from the PT tip (corresponding to the region where CFM was performed) were collected and used for the measurement of cell wall thickness. The sections were visualized in a CM100 TEM system (FEI, The Netherlands) using a Gatan Orius 1000 CCD camera (Gatan, Munich, Germany).

### Determination of Cell Wall Thickness

The cell wall thickness of wild-type and mutant PTs was determined from TEM cross sections described in subsection 1. Ten measurements around each section were averaged to give the cell wall thickness of an individual PT. All measurements were made with Fiji.

### Pollen Tube Diameter Measurements

To measure the diameter of fully turgid PTs, they were germinated and grown on silane-coated slides for 3 to 4 h. After carefully replacing the PTGM, the PTs were left growing for another 15 to 30 min before images were taken. Afterwards, plasmolysis was induced by replacing the growth medium with 15 % mannitol. Images were taken when full plasmolysis occurred, confirmed by a complete retraction of the protoplast from the tip of the PT. The diameter of turgid and plasmolyzed tubes, respectively, was measured 10 to 50 µm behind the tip using Fiji. The mean of five measurements within this range was taken as the diameter of an individual PT.

### CFM Measurements

The CFM analysis was performed as described in (16) and in (72). *Arabidopsis* PTs growing on silane-coated slides were visualized at a 400x magnification with DIC optics on an inverted microscope (IX 71, Olympus). MEMS-based microforce-sensors (FT–S540 and FT-S100) were used to measure the apparent stiffness and the viscoelastic relaxation of the PTs. Studies have indicated that the difference in anisotropic stresses at the tip are largely a function of the tip geometry, and even a constant isotropic elastic modulus along the pollen tube would yield different stiffness measurements at the tip compared to the shank (16). The indentation experiments were therefore performed 10 µm behind the apex and the apparent stiffness values were obtained by linearly fitting the resulting force-displacement curve. The maximal force was set to 4 µN, which resulted in an indentation depth of about 2 µm. For each PT, five measurements with four scans each were taken and *>*20 PT were analyzed per plant line. The resulting apparent stiffness data (Figure S3A) were used for FEM simulations. Additional parameters necessary for the analysis are listed in Figure S3 and Table 1. For the relaxation experiments, the PTs were also indented by the force probe at a distance 10 µm from the tip. The position of the indenter was kept for 5 s after the maximal force of 5 µN was reached. The force decay during the constant displacement was recoded. To determine whether the experimental setup contributes to the force relaxation, we measured a silicon cantilever (FS-C 15, SiMETRICS) that behaves in a purely elastic manner. In addition, we measured glass fibers, which have a geometry similar to the PTs, but a higher viscoelasticity. Data acquisition and control of the indenter were implemented in LabVIEW.

### Growth Rate Measurements

After the PTs had entered the microchannels, images were taken every 3 s during 20 min on a LM using an ORCA-D2 camera (Hamamatsu Photonics K.K., Hamamatsu). KymographClear, a macro toolset for Fiji was used to produce high-quality kymographs of the growing PTs, from which the growth rate was calculated using KymographDirect (73).

### FEM Simulation

The model is inspired by an earlier first-order model that estimated the elastic modulus and turgor pressure of the PT from CFM apparent stiffness measurements using Laplace’s law for thin-shell hollow tubes (16, 74, 75), given by

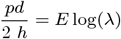

where *p* is the turgor pressure, *d* is the PT diameter, *h* is the cell wall thickness, *E* is the elastic modulus, and *λ* is the circumferential stretch.

The driving script for the Monte Carlo simulation in Abaqus was developed in Python.

The correlogram and PCA analysis plots were created in python using the sklearn and seaborn packages.

### Analytical Relation Describing the Apparent Stiffness for Indented Thin Shell Tubes

As described by Arnoldi and colleagues (76) and established for PTs by Burri and coworkers (72), the apparent stiffness *k* = *F/δ* of a pressurized cylinder measured under indentation can be described by

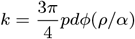

where *F* is the measured force, *δ* is the indentation depth, *p* is the turgor pressure, *d* is the PT diameter, and *φ*(*ρ/α*) is a geometric factor (72, 76). *φ*(*ρ/α*) can be further described by

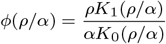

where *ρ* is the radius of the indenter, *Kn* are modified Bessel’s functions, and *α* is the cutoff distance given by

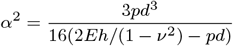

where *E* is the elastic modulus, *h* is the cell wall thickness, and *ν* is the Poisson’s ratio.

### Quantification and statistical analysis

Hypothesis testing for the measured apparent stiffness and wall thickness was performed using a one-way repeated-measures ANOVA omnibus test with the application of the Greenhouse-Geisser correction due to violation of the sphericity property. Pairwise comparisons were then made using multiple one-way paired t-tests. Significant differences in the diameter measurements and the simulated turgor pressure and elastic modulus values were tested using a one-way ANOVA omnibus test followed by the Games-Howell post-hoc test. All tests were performed with a 95 % confidence interval and the pairwise comparisons used the Holm-Bonferroni correction.

## ACKNOWLEDGMENTS

We are grateful to Markus Pauly (University of Dusseldorf) for providing xeg113–2 seeds, to Daniel Bollier (University of Zurich) for his ingenuity to improve the CFM, to Nick Jaensson (ETH Zurich, now at Eindhoven University of Technology) for valuable suggestions on optimizing the FEM/Monte Carlo model, and to Christina Westermann (University of Zurich) for comments on the manuscript. This work was supported by the University of Zurich, the ETH Zurich, and the Research and Technology Development Projects ‘Plant Growth in a Changing Environment’ (to U.G., B.J.N. and C.R.) and ‘MecanX – Understanding Physics of Plant Growth’ (to U.G., B.J.N., H.J.H. and C.R.), supported by SystemsX.ch, the Swiss Initiative in Systems Biology, and, in part, by an interdisciplinary grant of the Swiss National Science Foundation (CR22I2_166110 to U.G., B.J.N and H.J.H).

