## Supplemental Information for "Changes in cell wall biochemistry affect mechanical properties and growth rate of *Arabidopsis* pollen tubes"

**This PDF file includes:**

Supporting text
Figs. S1 to S3
Table S1
SI References

### Supporting Information Text

### FEM model details

Due to large intra-sample variance and skew in the experimentally determined input parameters, an elaborate Monte Carlo uncertainty quantification was performed within the FEM framework, starting with a parametric fit on all experimental observations. The turgid and plasmolysed PT diameters were best fit using a lognormal distribution due to a strong bias in their lower tails. This skewed distribution is expected, especially among mutant PTs, as smaller diameters indicate greater wall stress (1), which possibly resulted in the elevated frequency of PTs bursting before measurement. This is supported by previous studies where measurements of biological structures exhibiting growth have also been shown to follow the lognormal distribution (2–5). These fitted probability distributions were then sampled with Latin hypercube sampling (2), and the resulting samples were correlated using the Iman-Conover method (5). Input parameter realizations from these correlated distributions were subsequently simulated until convergence (minimum 100 realizations) for each mutant using the inverse FEM-based model, until the averaged PT elastic properties—turgor pressure and elastic modulus—could be extracted.

Since a large number of input realizations had to be simulated to obtain accurate statistics for turgor pressure and elastic modulus, the FEM-based model was set up to ensure robustness and computational efficiency, whilst not sacrificing on accuracy. The FEM mesh consisted of a cylindrical shell attached to a hemispherical tip with an imposed pressure boundary condition on the inside wall (total length: 100 µm), constrained by a pinned distal edge. The computational expenditure of the simulations were considerably reduced by cutting the PT in half and enforcing mirror boundary conditions. Since the ratio of PT diameter to cell wall thickness in *Arabidopsis* is usually less than 25, the shear stress between the inner and outer surfaces cannot be neglected, requiring the use of quadratic brick elements unlike in previous studies on lily PTs (6, 7). The PT cell wall material was characterized using the Saint Venant-Kirchhoff hyperelastic material model, in which the Poisson ratios were set to 0.2 after a sensitivity analysis had shown that varying it in the range between 0.0 and 0.4 had a negligible effect on the apparent stiffness. PT cell walls have also been hypothesized to possess anisotropic properties due to the preferential orientation of cellulose microfibrils (8). However, a previous CFM study (7) concluded that the PT cell wall possesses only a very slight degree of anisotropy, which is largely masked by the effects of geometry and turgor pressure. Based on this conclusion, the current model neglects the effect of anisotropic material properties, as the geometry and range of applied turgor pressure remain unchanged. The contact indentation between the PT and the rigid microelectromechanical systems (MEMS)-based force sensor probe was considered frictionless with finite sliding. It was simulated using a surface-to-surface discretization with constraints enforced by Lagrange multipliers, along with appropriate damping to stabilize rigid body modes.

### Analytical relation details

To circumvent this constraint in estimating the relaxation modulus, we used an analytical relation developed for the nano-indentation of thin shell tubes in combination with the turgor pressure obtained from the FEM simulations. This relation is given by

$$F = \frac{3\pi}{4}\delta p d \phi \qquad [1]$$

where $F$ is the measured force, $\delta$ is the indentation depth, $p$ is the turgor pressure, $d$ is the PT diameter, and $\phi$ is a factor based on the relaxation modulus $E^r$ and the cell wall thickness $h$ (9, 10) (see STAR Methods). This relation allows for the force $F$ to be transformed into the relaxation modulus $E^r$.

Constitutive models of biological tissues often use the generalized Maxwell linear viscoelastic model to capture relaxing systems (11, 12). This model consists of a viscous damper and an elastic spring in series, the combination of which are placed in parallel with another elastic spring to describe the stress-strain behavior of a material. The relaxation modulus $E^r$ for such a system is given by

$$E^r(t) = E^\infty + \sum_i E_i^0 \exp^{t/\tau_i} \qquad [2]$$

where $E^\infty$ is the equilibrium modulus at 5 s, representing the purely elastic behavior, $E^0$ is the component of the elastic modulus $E$ subjected to time-dependent behavior, and $\tau = \eta/E^0$ is the relaxation time, where $\eta$ is the viscosity.

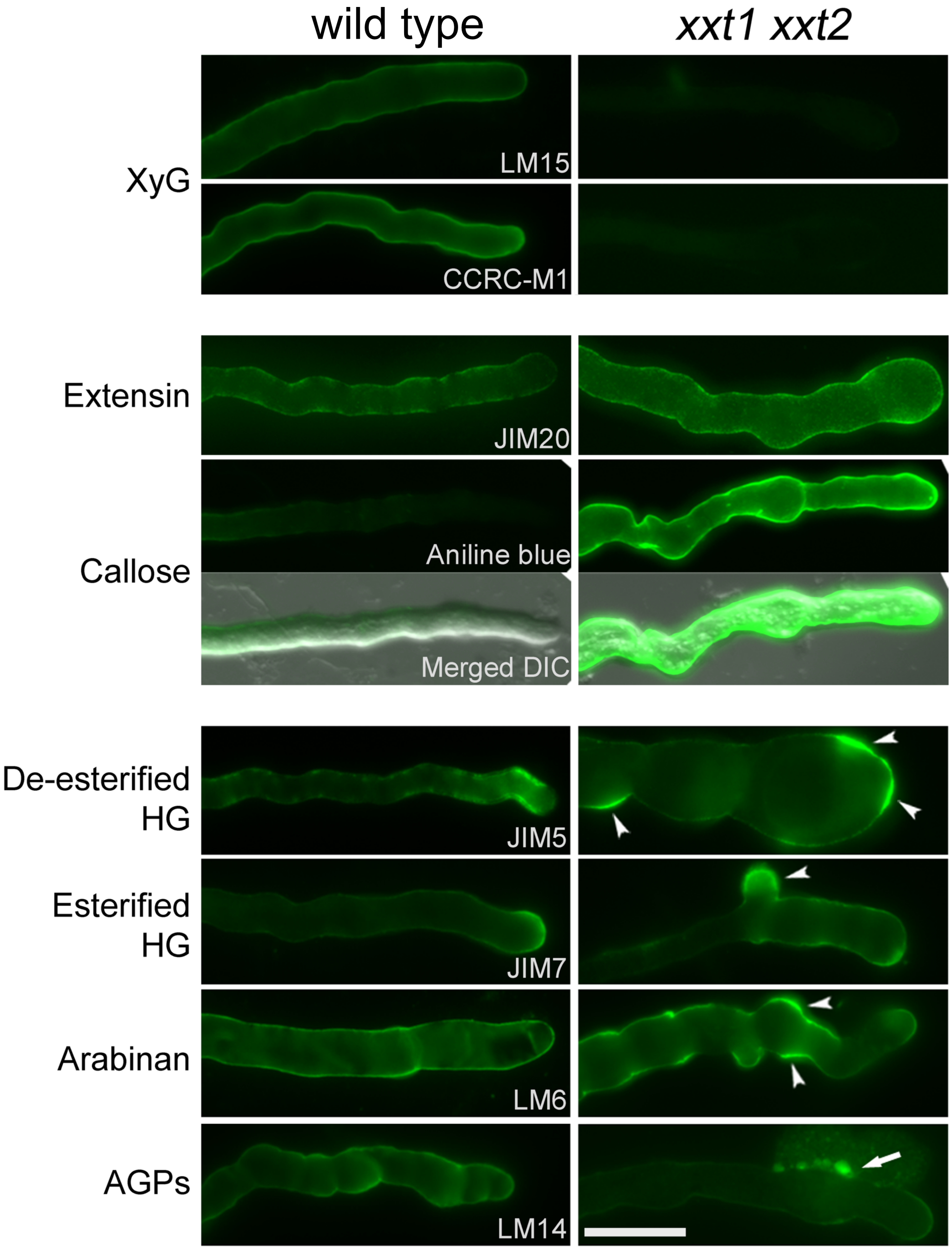


**Fig. S1. Immunodetection with Fluorescently Labelled Antibodies Against Cell Wall Epitopes Reveals Cell Wall Defects in *xxt1 xxt2* Pollen Tubes.**
Immunodection with LM15 and CCRC-M1 antibodies against xyloglucan shows no labelling in *xxt1 xxt2* PTs, staining with JIM20 antibodies against extensins and aniline blue for callose shows stronger signals in *xxt1 xxt2* compared to wild type PTs. Aberrant staining patterns are seen with JIM5 and JIM7 antibodies against methyl-esterified and de-esterified homogalacturonan, respectively, LM6 antibody against the arabinan side chain of rhamnogalacturonan I, and LM14 antibody against arabinogalactan proteins. Arrowheads/arrow indicate aberrant structures in mutant *xxt1 xxt2* PTs. Scale bars: $20\mu m$.

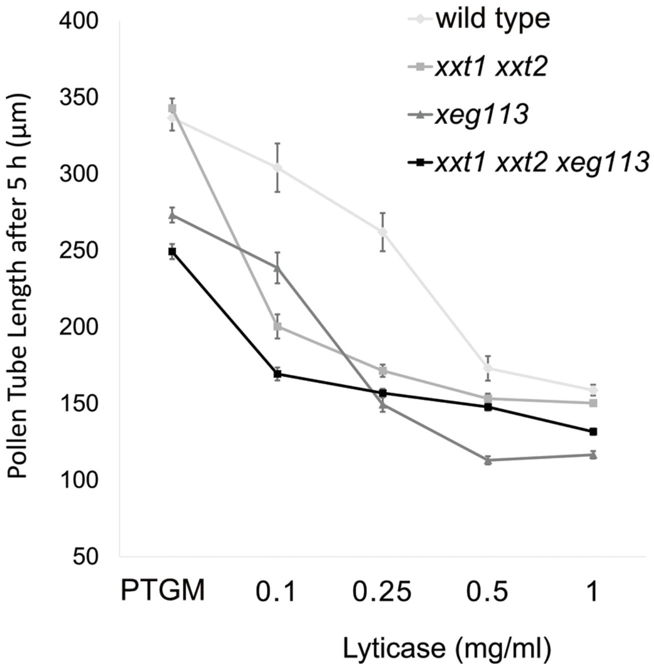


**Fig. S2. Effect of Lyticase Treatment on Wild Type and Mutant Pollen Tube Growth.**
Increasing concentration of lyticase in pollen germination medium reveals the susceptibility of the different mutants to callose digestion compared to the wild type: growth of *xxt1 xxt2* and *xxt1 xxt2 xeg113* PTs shows the highest sensitivity to lyticase, with a strong reduction of the growth rate at $1\,\mathrm{mg/mL}$ lyticase.

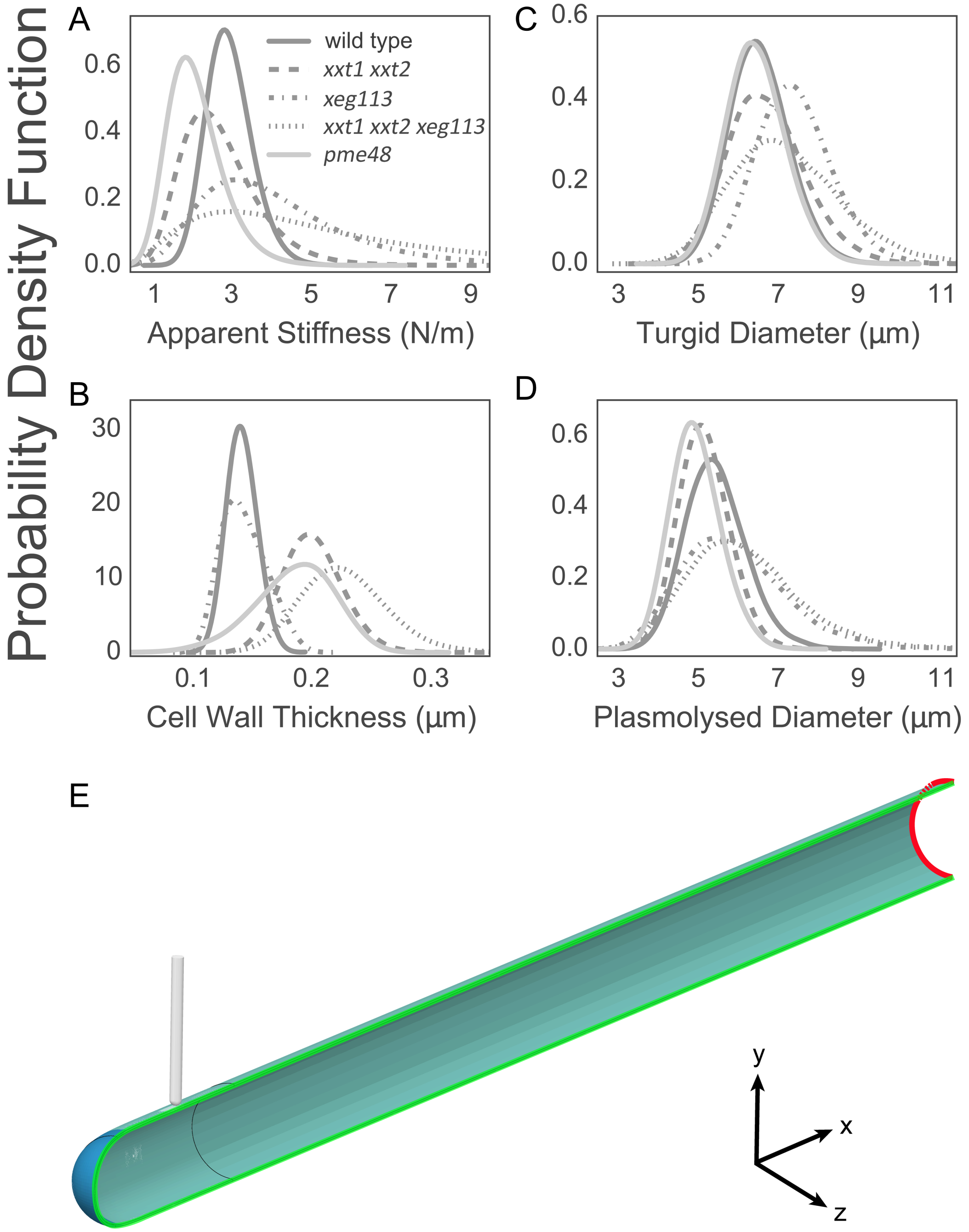


**Fig. S3. Mechanical Properties of Wild Type and Mutant Pollen Tubes.**
(A–D) Probability density functions of input parameters for the FEM modelling with Monte Carlo uncertainty quantification, including CFM-determined apparent stiffness (A), TEM-determined cell wall thickness (B), and the diameter of turgid (C) and plasmolysed PTs (D).
(E) The FEM model of the fully turgid pollen tube (with length of $100\,\mu\mathrm{m}$) before indentation. The boundary conditions during inflation and indentation are illustrated in red (symmetry about the YZ-plane), and green (symmetry about the XY-plane).

**Table S1. Quantitative Phenotypic Data**

Phenotypes are expressed as a percentage of all seeds sown on MS plates. WT_*pme48* denotes PTs of wild type plants segregating from the *pme48* mutant line. Phenotyping of the *pme48* mutant PTs was done separately from the other mutant lines.

| | wild type | *xxt1 xxt2* | *xeg113* | *xxt1 xxt2 xeg113* | *pme48* | WT_*pme48* |
|---|---|---|---|---|---|---|
| Germination | 21 | 30 | 23 | 32 | 51.4 | 59.2 |
| Bursting | 2.9 | 11.9 | 21.7 | 42.6 | 8.8 | 0.4 |
| Swelling | 0.6 | 5.5 | 1.9 | 25.1 | 24.4 | 2.2 |
| Branching | 0.6 | 1.2 | 1.4 | 2.7 | 26.9 | 0.2 |
| Total Deformed | 4.4 | 18.6 | 25.1 | 70.4 | 60.1 | 2.8 |
